# Advancing Aquaculture Monitoring through Self-Powered Remote Systems: Piezoelectric Energy Harvesting Approach

**DOI:** 10.1101/2023.12.31.573301

**Authors:** Iman Mehdipour, Luca Fachechi, Francesco Rizzi, Massimo De Vittorio

## Abstract

This paper introduces the conceptualization and implementation of a sea wave piezoelectric energy harvesting system designed for a self-powered IoT buoy sensor, tailored for the remote monitoring of water quality variables in fish farming. The developed autonomous system undertakes periodic measurements of temperature, pH, and dissolved oxygen, with the gathered data transmitted locally through a low-power wide-area network protocol to a gateway. The gateway is intricately connected to a cloud service, facilitating efficient data storage and visualization. At the heart of this innovation lies a novel buoy design capable of harnessing energy from sea surface waves. The proposed piezoelectric energy harvesting system comprises two pivotal components: an energy transfer mechanism responsible for capturing incident sea surface wave energy and a piezoelectric fin serving as the transducer. Extensive data analysis of sea wave patterns spanning nearly 25 years informs the design process. Subsequently, a prototype is meticulously crafted, fabricated, and rigorously tested in a controlled environment, showcasing promising results. The outcome is a self-powered remote monitoring tool poised to revolutionize fish farming scenarios by enabling seamless, automatic data acquisition, and storage.

## 1. Introduction

The meticulous supervision of water-quality parameters, encompassing critical facets such as dissolved oxygen, temperature, and pH, assumes a position of paramount importance within the realm of pisciculture. Traditionally, practitioners in pisciculture have been tasked with the manual collection of data pertaining to these variables, their measurement frequencies intricately entwined with the fluid dynamics of the pond and its ambient milieu [1-3]. However, this manual approach is burdened by temporal demands, exceedingly low acquisition rates, and the imperative for frequent recalibration. In response to these constraints, a discernible impetus has emerged, steering the trajectory toward the development of automated remote monitoring devices. These technological innovations aspire to alleviate the data collection burden on fish farmers while concurrently elevating the frequency of data acquisition [4-6]. For example, a recent battery-powered system measures temperature, pH, and dissolved oxygen, transmitting data locally through a low-power wide-area network to a cloud service. This novel buoy design, constructed from readily available materials, not only lends stability to the Internet of Things (IoT) device and immersed probes but also showcases promising results as a low-cost remote monitoring tool for automatic data acquisition and storage in fish farming scenarios [7].

In the broader context of marine environments, where diverse mechanical energy sources such as waves, tides, and currents abound, substantial research has been dedicated to technologies for energy conversion [8]. The vast, predictable, and reliable energy reservoir of the ocean has sparked a surge in interest in electricity generation, with piezoelectric materials standing out for their threefold higher energy generation density among small-scale harvesters. These materials offer simplicity in attachment, lack of moving parts, and freedom from frequent maintenance, providing advantages like direct energy-to-electricity conversion, compactness, structural simplicity, cleanliness, light weight, stability, and high sensitivity to small strains. While piezoelectric harvesters may produce relatively modest electricity, it suffices for low-energy consumption electronics, particularly oceanic sensors.

A recent study delves into the imperative of battery elimination in oceanic sensors, exploring various direct energy harvesting methods with a specific focus on piezoelectric energy harvesters. Presenting an atlas of 85 recently reported designs, the research categorizes them based on configurations and provides comprehensive insights into materials, coupling modes, locations, and power ranges. It critically discusses concepts and addresses challenges in the field [9]. Additionally, the potential of ocean wave energy as a renewable resource is examined, with a particular emphasis on piezoelectric energy harvesters (PEH) as a means to self-power ocean monitoring sensors [10].

Categorizing PEH into direct-coupling, frequency-increasing, flow-induced vibration, and multi-mechanism composite types, the study provides detailed analyses and insights into their materials, principles, applications, and shortcomings. The aim is to guide future research in efficiently converting ocean wave energy into electrical energy for sensor deployment in the ocean environment.

The integration of a multiphysics system is developed for a one-way plucking-driven piezoelectric wave energy harvester (OPDPWEH) with a floating cylindrical buoy, a frequency up-conversion mechanism, and piezoelectric bulk composite cantilever beams [11]. A sequential rotary-driven piezoelectric wave energy harvester (SRD-PWEH) with a cylindrical buoy is investigated by a sequential-drive rotating mechanism using one-way bearings and a piezoelectric harvesting component based on a circumferentially arranged array of piezoelectric composite cantilever beams [12]. Moreover, a Flexible PiezoElectric Device (FPED) is designed to provide a continuous and stable energy supply for Fish Aggregating Devices (FADs) and islands, utilizing FPEDs coated with piezoelectric paint [13]. Experimental, theoretical, and computational approaches are employed to assess FPED characteristics, wave interactions, and real-sea performance, demonstrating the effectiveness of the FPED under extreme conditions and validating the theoretical and computational models as alternative tools for FPED working parameter selections. An embedded patch-based energy harvesting approach on a marine composite boat [14] is proposed by employing piezoceramics (PZT) and ZnO as patch materials and evaluates the performance of the harvester exposed to wave-induced loads. Furthermore, a novel approach to wave energy harvesting is explored with the proposal of a Cylindrical-Conical Buoy Structure and Magnetic Coupling-based Piezoelectric Wave Energy Harvester (C-PWEH) [15]. In this approach, utilizing the buoy’s oscillations driven by waves and magnetic coupling to deform piezoelectric patches, the device optimizes wave energy conversion efficiency.

The central focus of this paper lies in energy harvesting from sea surface waves through piezoelectric energy convergence to power Internet of Things (IoT) sensors in aquaculture applications. The primary objective is to extend the battery life of the IoT device through a piezoelectric energy harvesting system, comprising two main components: an energy transfer mechanism from the surface to the deep water and a piezoelectric transduction mechanism. The subsequent sections of this document are meticulously organized. Initially, the design process for an optimal piezoelectric energy harvester is expounded upon. Subsequently, each component of this process, encompassing the analysis of ambient source data, the selection of piezoelectric materials, the design of the energy transfer and transduction mechanism, and the design or selection of the power management system, is discussed in intricate detail. Finally, the last section succinctly concludes the comprehensive work undertaken in this study.

## 2. Design parameters for piezoelectric energy harvesting systems

The presented schematic intricately outlines a closed-loop process meticulously crafted to achieve the epitome of design excellence for an energy harvester, thereby maximizing its performance potential. Within this algorithmic framework, the ambient energy source functions as the primary input, positioned as an entrance gate, while the remaining algorithmic components orbit as integral elements of a feedback loop. Each of these components assumes a pivotal role in optimizing the harvester’s performance.

The paramount importance lies in the judicious selection of the piezoelectric material, serving as the primary parameter within the transduction mechanism. Simultaneously, the power management system, characterized by minimal energy losses and a lower threshold voltage, represents another indispensable parameter within this loop, demanding meticulous optimization. Moreover, the power consumption of the sensor and the telecommunication protocol stands as fundamental input parameters essential for the design of any energy harvesting system. These parameters must be tailored to meet the minimum requirements dictated by the data acquisition or monitoring strategy.

Subsequently, propelled by considerations encompassing the potential energy of the ambient source, the energy conversion efficiency of the piezoelectric material, the performance characteristics of the designed transducer, and the power losses inherent to the power management circuit, decisions will be made regarding the design of the energy harvesting system. These decisions will be guided either towards creating a fully self-powered IoT sensor or extending the battery life of a remote sensor.

Figure 1 vividly illustrates the design process for an optimal piezoelectric energy harvester through a Sankey diagram, delineating the generated and lost energy at each design stage. The diagram reveals that a substantial portion of energy dissipates at the initial stage due to the piezoelectric conversion mechanism and mechanical coupling. Additionally, electrical leakage accounts for another form of loss in the power management system, ranging from 20 to 30 percent. Lastly, voltage regulation loss occurs during the adjustment of the threshold voltage for consumption in sensors and telecommunication modules, constituting around 20 to 25 percent.

**Figure 1.**
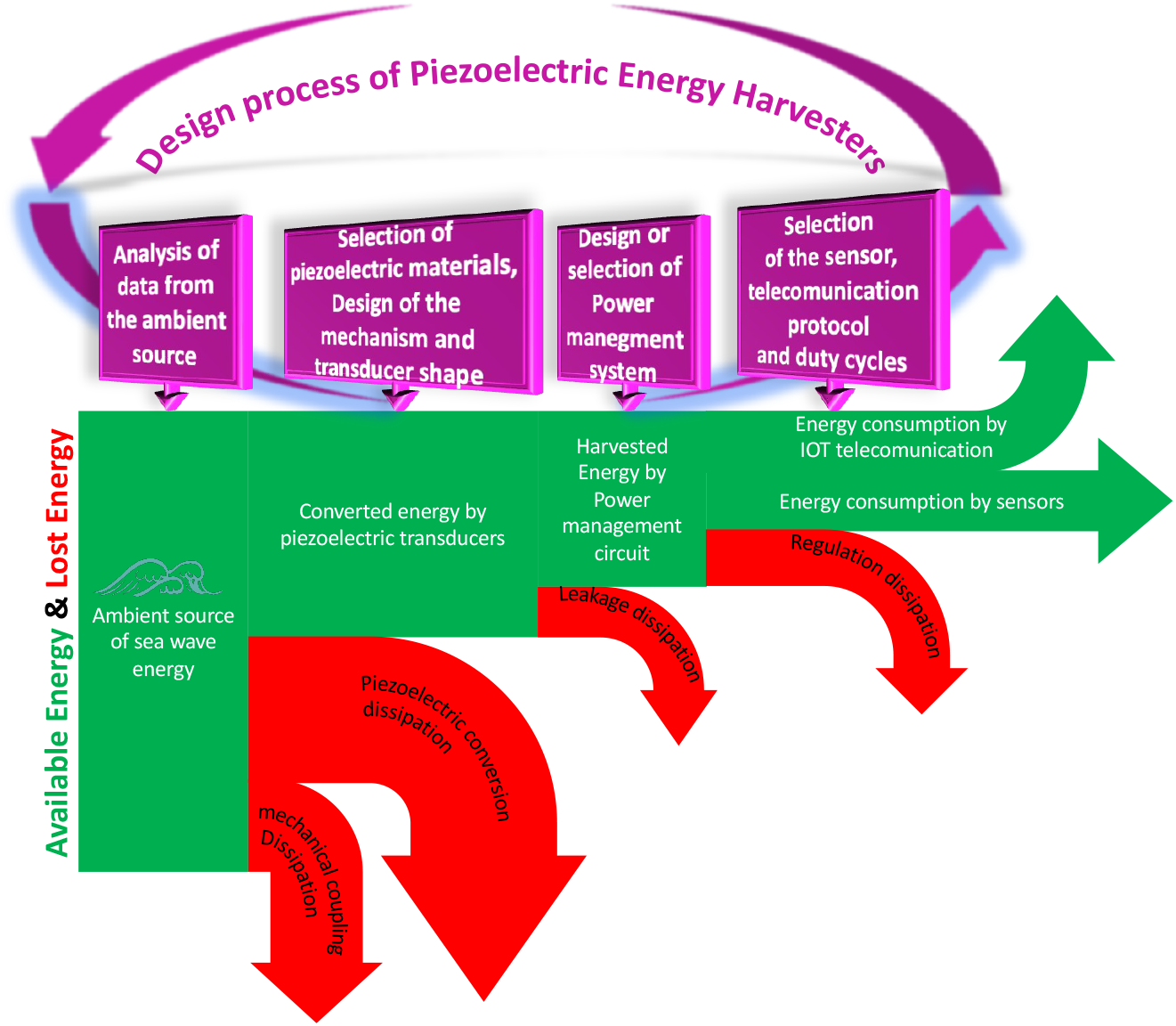
Design process for an Optimal Piezoelectric Energy Harvester by considering Sankey diagram of the generated and lost energy in each design stage.

In essence, the primary objective of this flowchart detailing the design process for an optimal piezoelectric energy harvester is to minimize losses and enhance performance at each stage. This is achieved through meticulous mechanical design, judicious material selection, circuit design with high-performance components, and the selection of low-power consumption for sensors and telecommunication protocols, coupled with an appropriate duty cycle for measurements and data transfer.

In the subsequent subsection, we delve into a comprehensive exploration of three fundamental design parameters: the ambient energy source, the selection of an appropriate piezoelectric material, and the meticulous design of the transduction mechanism. These aspects are scrutinized in detail with the objective of crafting a piezoelectric energy harvester tailored for application in the advancement of aquaculture monitoring through self-powered remote systems.

### 2.1. Ambient source (Wave spectrume)

The efficacy of harnessing piezoelectric energy from an ambient source hinges upon a multitude of critical factors. These encompass the amplitude and frequency of oscillations, their directionality, stability, and duration, all of which exert a profound influence on both energy output and system reliability. Equally pivotal is the assessable harvestable area and its compatibility with the piezoelectric harvester, both of which assume paramount importance in practical deployment scenarios.

Environmental conditions, energy requisites, and the efficiency of the conversion process must be judiciously evaluated, alongside considerations such as scalability, maintenance, and cost-effectiveness. Adapting the energy harvesting system to harmonize with these multifaceted features constitutes an imperative step toward optimizing performance, thereby harnessing mechanical vibrations and movements as a sustainable and reliable power source.

Thanks to satellite technology facilitating data collection, a comprehensive dataset spanning a quarter-century has been acquired for the location of Mattinata in the Adriatic Sea, in Figure 2, utilizing sea data as the ambient source.

**Figure 2.**
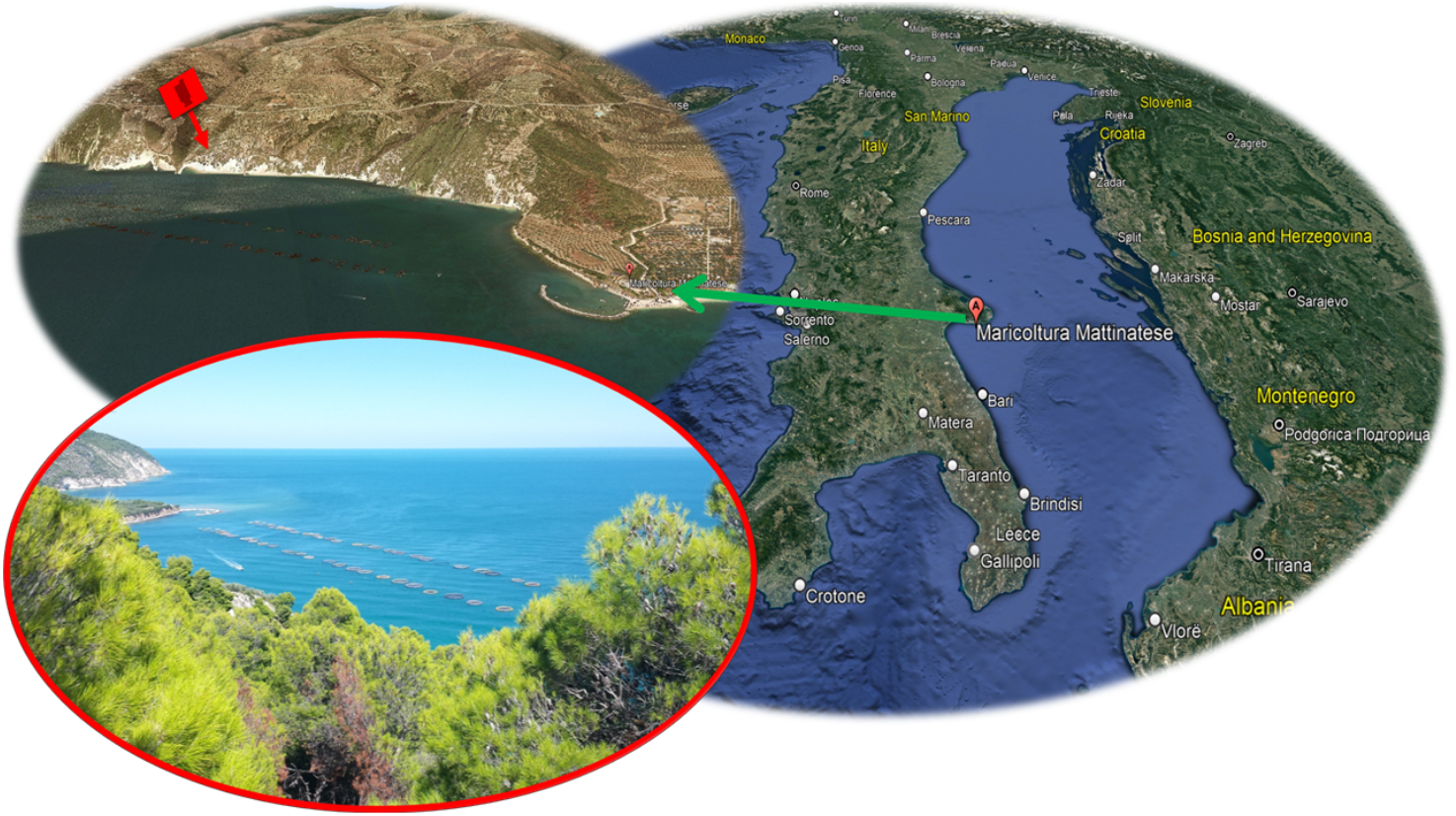
Maricoltura Mattinatese (41.725804, 16.109511) fish farm station, Mattinata, Province of Foggia, Italy

This dataset encompasses a wide array of parameters, including temperature, water current in three dimensions extending from the sea surface to various depths, as well as wave height and period at the sea’s surface.

In the course of this investigation, the design of the piezoelectric energy harvesting transduction mechanism is rooted in the data obtained from the sea surface, specifically focusing on the wave height and period.

The sea surface wave height and period data have been sampled at an hourly rate spanning from 1993 to 2021, encompassing the sea’s free surface. The time history of sea surface wave height, vividly showcasing instances where wave heights can surge to an impressive 4 meters during rare occurrences. To ensure the robust design of the energy transfer mechanism and harvester transducer, it is imperative to base calculations on the most probable sea wave height.

The Temporal Profile of Sea Surface Wave Period illustrates variations spanning from 2 to 12 seconds. A crucial parameter for the design of the piezoelectric transducer is the frequency of the external force, with the sea wave frequency serving as the excitation frequency in this project. Consequently, a longer sea wave period translates to a lower excitation frequency for the transducer. As mentioned earlier, it is imperative to account for the most probable sea wave period when designing both the energy transfer mechanism and the harvester transducer, a topic we delve into in the following section.

In the ensuing figures, probability functions have been generated to highlight two critical parameters in the design of a mechanism for energy harvesting from sea surface waves, namely, surface wave period and wave height. Figure 3 unequivocally illustrates that the most probable surface wave height is approximately 20 cm. Furthermore, the most likely wave period hovers around 2 seconds, equivalent to an approximate frequency of 0. 5 Hz.

**Figure 3.**
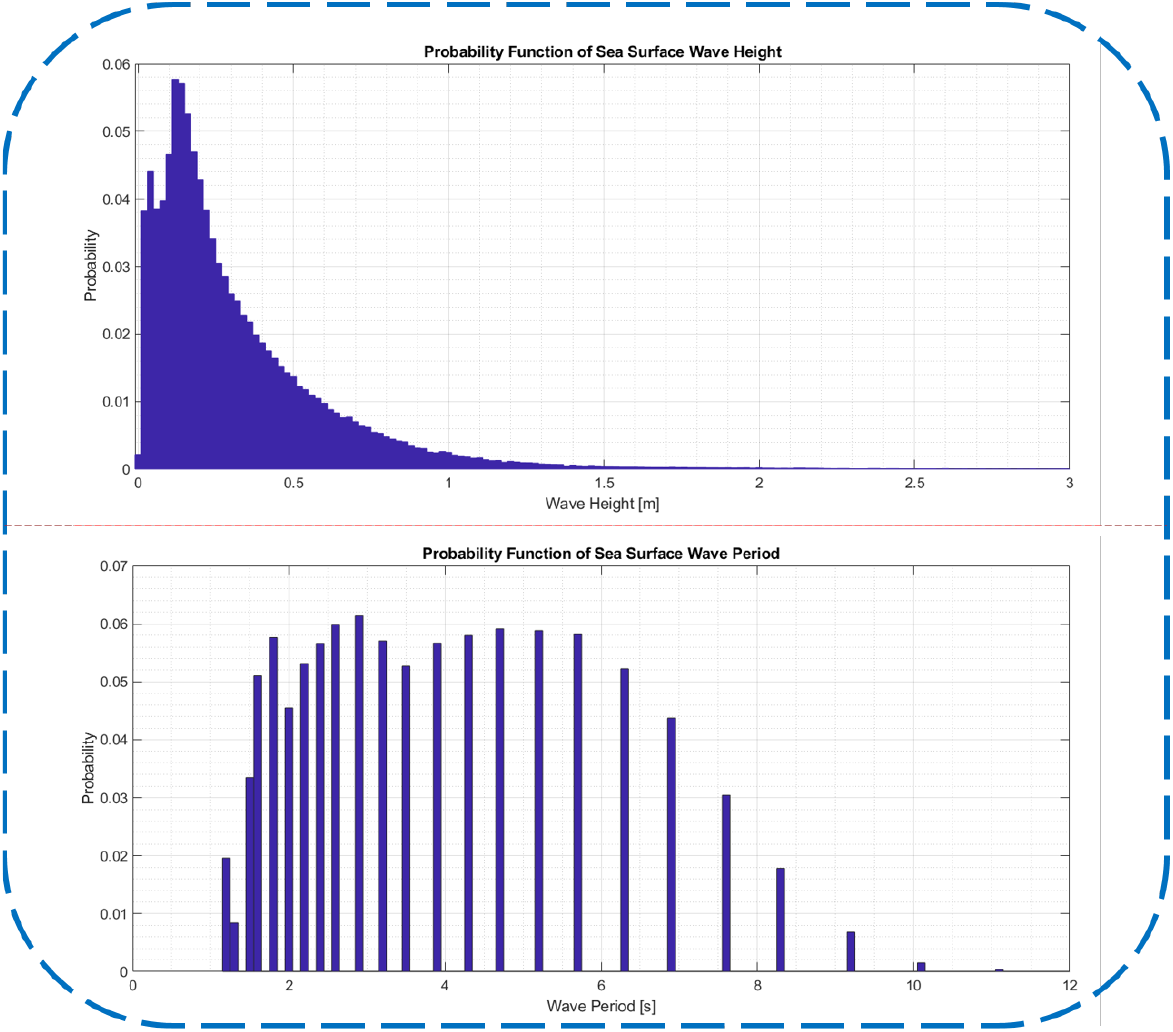
Probability density functions of sea wave height and period for a time slot from 1993 to 2021 in Maricoltura Mattinatese (41.725804, 16.109511).

Sea waves play a crucial role in supplying input energy to piezoelectric energy harvesters. In Figure 4, we present a comprehensive view of wave statistics, showcasing the joint probability distribution of significant wave height and wave peak period for the proposed fish farm. The background colour in the figure serves as a visual representation of the probability of occurrence for each wave condition, expressed as a percentage. This percentage is derived by dividing the occurrences of each wave condition by the total number of occurrences.

**Figure 4.**
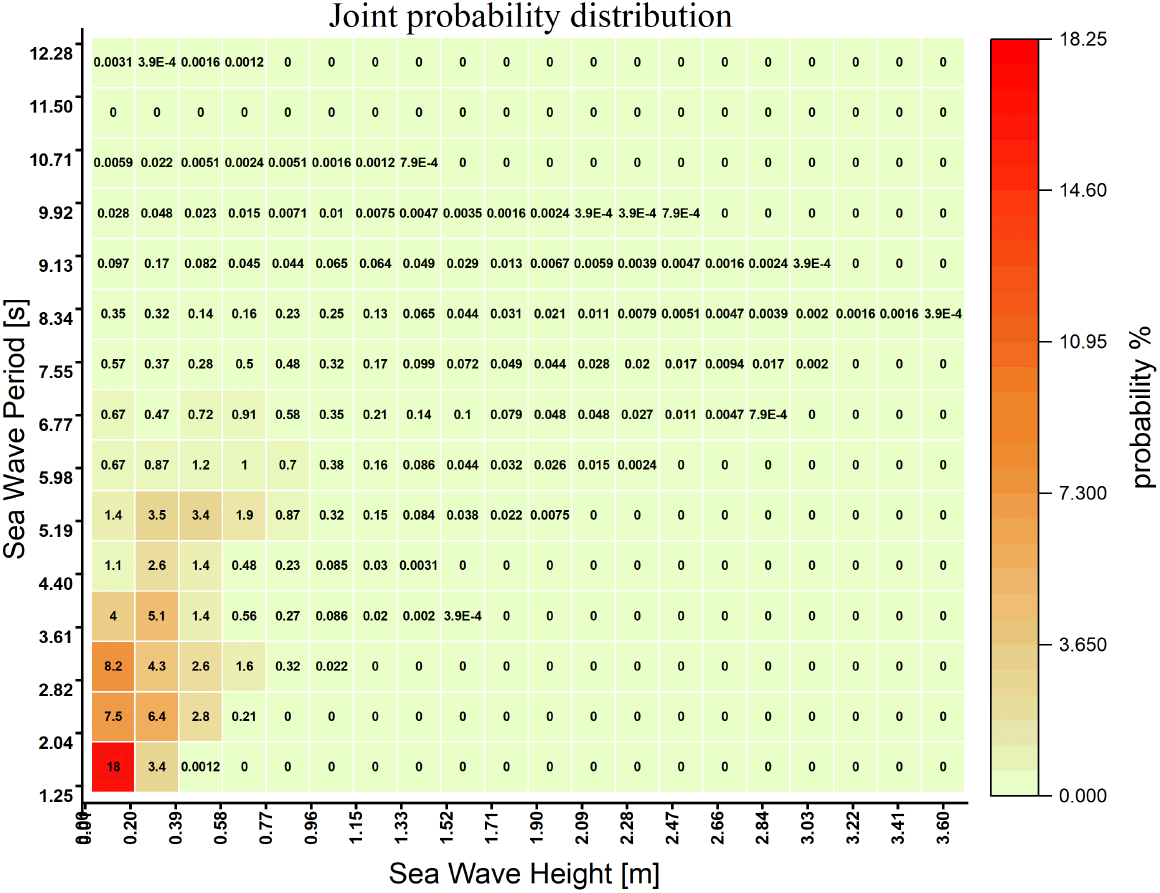
Joint probability distribution at significant wave height and period.

Analyzing the wave field data in Figure 3, we observe that waves with a significant height of ≤ 1 m and a period of ≤ 7s dominate, constituting a substantial 91.33% of the total occurrences. Notably, the highest probability, accounting for 18%, is associated with significant wave heights ranging from 0.1 to 0.2 m, peak periods between 1.2 and 2s, and an average peak period falling within the 2 to 4s range, with significant wave heights spanning from 0.1 to 0.4 m. This detailed breakdown provides valuable insights into the prevailing wave conditions and their probabilities, essential for optimizing the design of the proposed fish farm and its associated energy harvesting systems.

The wave power level (*P*) per unit width in a wave or wave energy flux (kWh/m) for irregular waves can be calculated based on the peak period (*T*_*e*_) and significant wave height. Here’s the breakdown [16-18]:

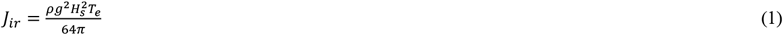

The energy period (*T*_*e*_) is linked to the peak wave period (*T*_*p*_) through *T*_*e*_ = *α T*_*p*_, with a coefficient (*α*) set to 1 for this study [18-20]. The scatter power diagram in Figure 5 visually represents the occurrence of wave power levels at different significant wave heights and peak periods. The probability of occurrence for the peak period relative to significant wave height is expressed as a percentage, with intervals of 1s and 0.2m, respectively. The wave power level is quantified in kWh/m and depicted using a color scale. Upon analysis, it’s evident that waves with maximum power potential (≥8 kWh/m) exhibit peak periods between 9s and 13s, coupled with significant wave heights ranging from 2.1m to 2.7m. This specific range represents only 0.15265% of the total share, emphasizing the unique conditions required for optimal wave power generation.

**Figure 5.**
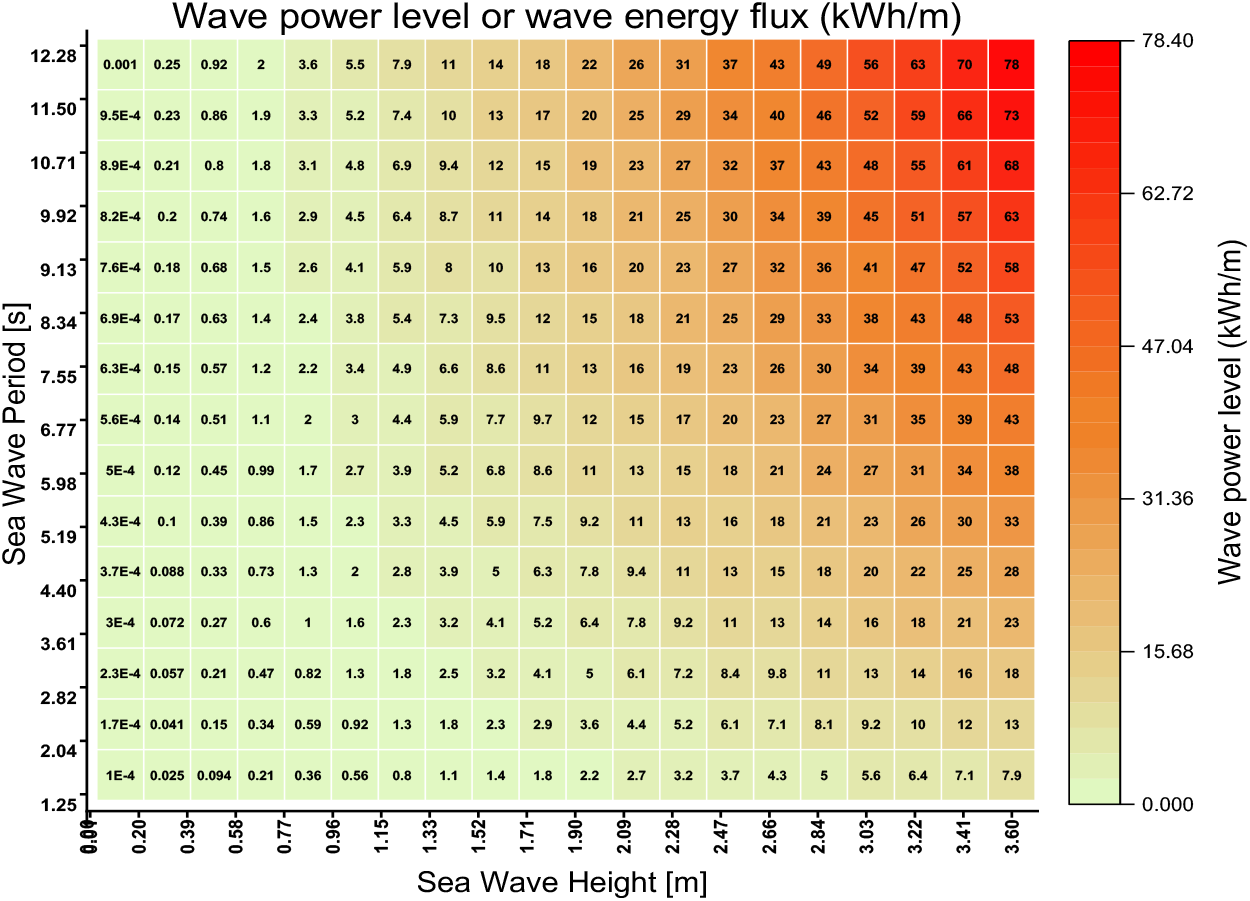
Wave power level (kWh/m) occurrence at significant wave height and period.

As a consequence, our principal objective is to engineer a mechanism capable of oscillating at the most probable frequency, which is 0.25 Hz, functioning as the resonance frequency. Furthermore, this mechanism aims to convert the surface wave height into an excitation displacement for the piezoelectric transducer located at a deeper water level.

The examination of normalized wave power levels, coupled with a joint probability distribution, stands as another statistical dimension under scrutiny in this investigation. As illustrated in Figure 6, this heatmap is a product of the joint probability distribution from Figure 4 and the wave power levels from Figure 5. This particular parameter unveils the most likely and elevated wave power levels primed for energy harvesting, serving as a valuable guide for refining and optimizing our harvester to achieve peak performance within this specific range. Figure 6 eloquently illustrates that the pinnacle of both probability and wave power levels is situated in the wave height spectrum ranging from 40 cm to 1 m, corresponding to wave periods spanning 5 to 9 seconds.

**Figure 6.**
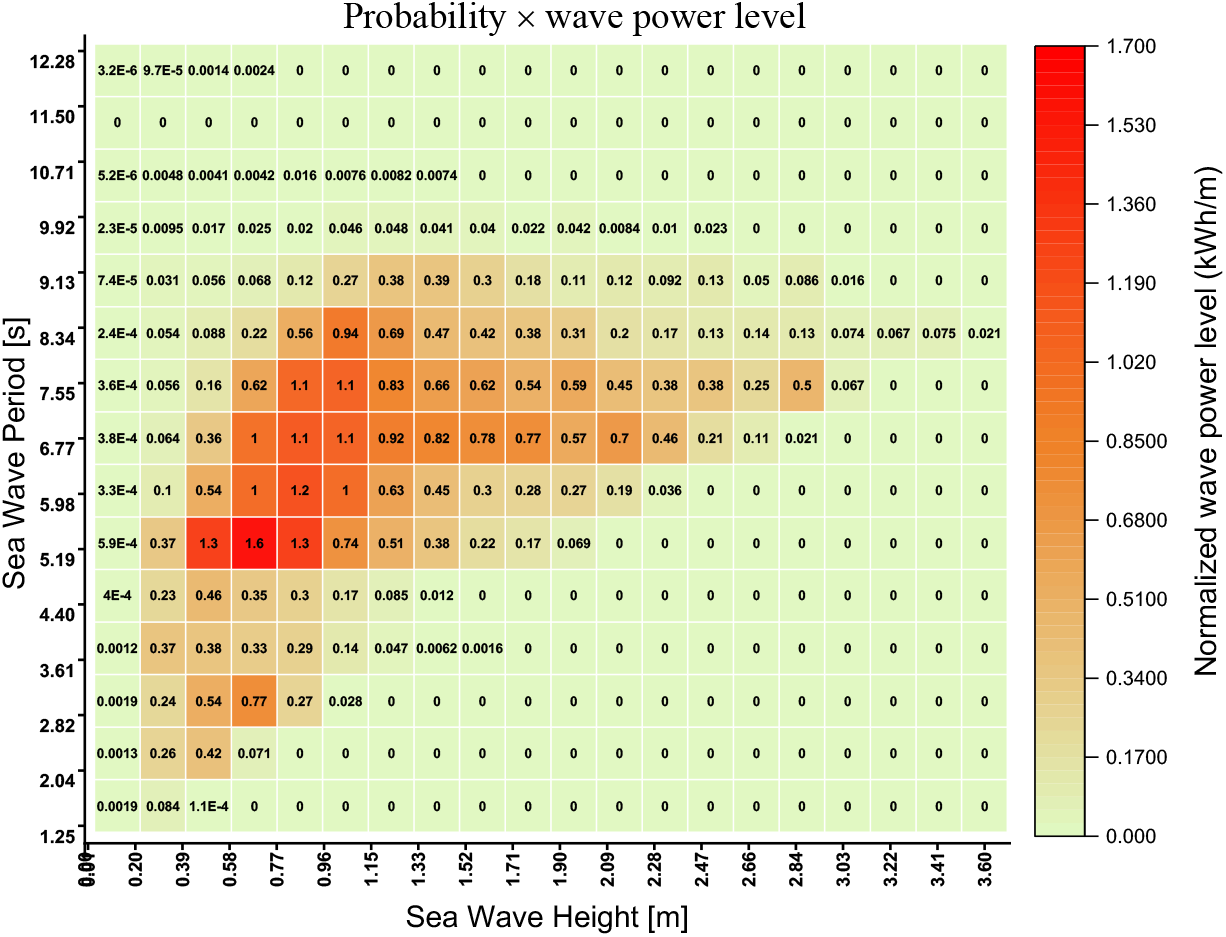
Normalized wave power level (kWh/m) occurrence at significant wave height and period according to joint probability distribution.

### 2.2. Piezoelectric materials

PZT, or Lead Zirconate Titanate, PolyVinylidene DiFluoride (PVDF), and Macro-Fiber Composite (MFC) stand out as primary piezoelectric materials for extracting electrical energy from the oceans [9]. PZT takes the lead in ocean energy harvesting due to its exceptional dielectric and piezoelectric properties, along with a superior d31 value. Its high Curie temperature, marking the point at which piezoelectric materials lose their piezoelectricity, enhances its versatility. Regular chemical modifications over time have broadened PZT’s applicability, maintaining its status as the most extensive family of piezoelectric materials, offering diverse properties at a cost-effective rate. Despite its brittleness and minimal elongation at break (0.1%), making it less flexible, PZT remains the preferred choice for energy harvesting due to its outstanding properties and affordability.

On the contrary, PVDF, known for its flexibility, finds application in scenarios requiring higher load-bearing capabilities. With a significantly lower Young’s modulus compared to PZT, PVDF exhibits a 10% elongation at break. However, achieving higher voltage amplitude in PVDF requires a higher percentage of elongation compared to PZT materials with the same input mechanical energy. Despite its flexibility, PVDF’s power density is much lower than that of PZT, presenting challenges in applications demanding high output power. Given the substantial mechanical energy in the sea, coupled with limited mechanical displacement to induce high elongation in PVDF, neither PZT nor PVDF proves suitable for this application alone. Composite piezoelectric materials like MFC offer a trade-off, providing both higher flexibility and power density than PZT and PVDF, respectively. Considering these factors, it is strongly recommended to explore composite piezoelectric materials in future designs for their favorable properties, including a lower Young’s modulus than PZT and a higher *d*_*31*_ constant than PVDF.

In this study, DuraAct patch transducers, flexible piezoceramic plates or films embedded in a polymer along with their contacts, are employed. This design not only mechanically preloads the brittle ceramic but also provides electrical insulation, making it a suitable option for sea wave energy harvesting systems. The compact design, inclusive of insulation, enhances user handling, and the possibility of embedding the patch transducer in a composite material adds to its versatility.

In this study, we’ve bimorphed each fin using two patches strategically positioned on the fin’s top and bottom. For a detailed breakdown of the DuraAct Patch P-876.A12’s geometrical and material properties, refer to Table 1. The piezoelectric material employed, identified as soft PZT with the commercial moniker PIC255, is thoroughly outlined in Table 1 and its specific material properties are neatly organized in Table 2.

**Table 1.**
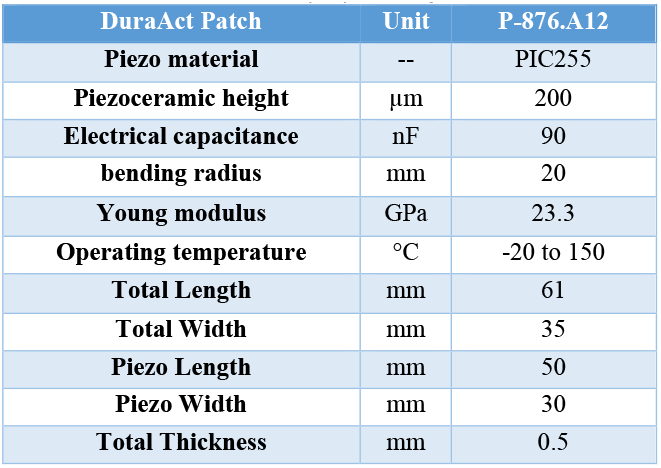
Geometrical and material properties of DuraAct Patch P-876.A12.

**Table 2.**
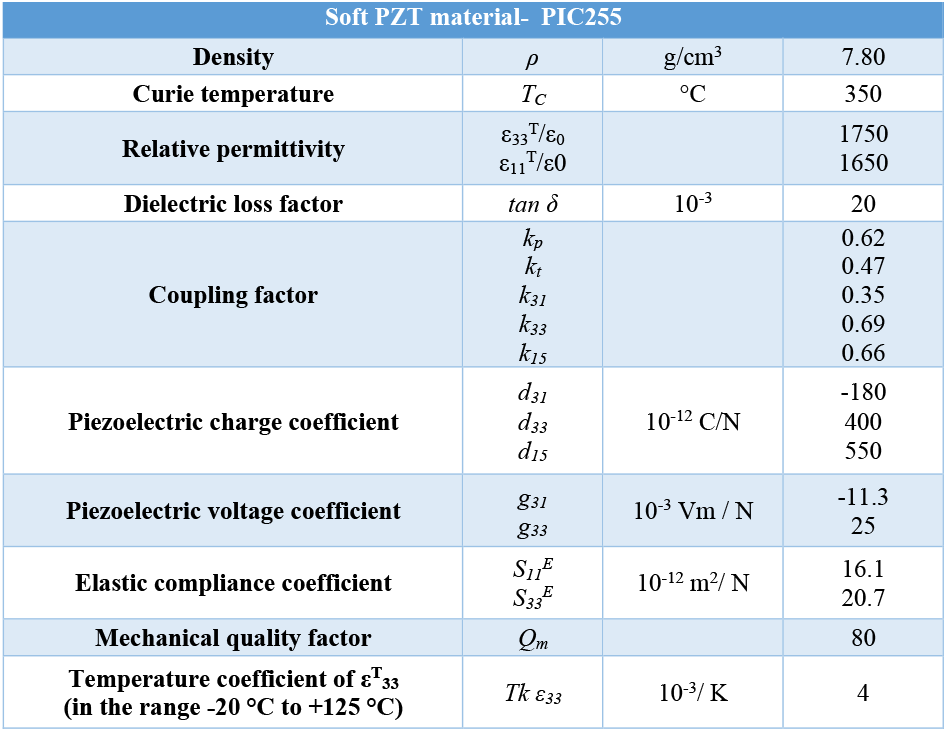
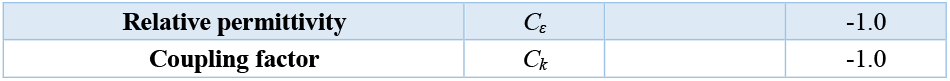
Piezoelectric material (PIC255) properties of DuraAct Patch P-876.A12.

Given the DuraAct Patch P-876.A12’s high Young’s modulus, notably standing at 23.3 GPa as indicated in Table 1, it becomes imperative for the substrate stiffness to surpass this value. This ensures the efficient transfer of the excitation force throughout the fin, particularly to the clamped section where both DuraAct patches find their place. Considering the unique application of this setup in water for harvesting energy from sea waves, we propose the use of a carbon fiber sheet embedded in an epoxy resin matrix. Table 3 provides a concise overview of the properties of this suggested material. Noteworthy attributes include a thickness of 0.5 mm and a transversal Young’s modulus of 70 GPa, making it a fitting and reliable choice for the demands of this application.

**Table 3.** Properties for Carbon/Epoxy Composite Sheet.

| Carbon fiber- Epoxy resin matrix | Unit | value |
| --- | --- | --- |
| Thickness | mm | 0.5 |
| Coefficient of thermal expansion - Longitudinal | $\times 10^{-6} \text{ K}^{-1}$ | 2.1 |
| Coefficient of thermal expansion - Transverse | $\times 10^{-6} \text{ K}^{-1}$ | 2.1 |
| Compressive Strength - Longitudinal | MPa | 570 |
| Compressive Strength – Transverse | MPa | 570 |
| Density | $\text{g cm}^{-3}$ | 1.6 |
| Shear modulus - in-plane | GPa | 5 |
| Shear strength - in-plane | MPa | 90 |
| Volume fraction of fibres | % | 50 |
| Young's Modulus - Longitudinal | GPa | 70 |
| Young's Modulus - Transverse | GPa | 70 |

### 2.3. Design of the transducer and energy transfer mechanism

This section is divided into two key segments: the design of the energy transfer mechanism and the configuration of the piezoelectric energy harvester transducer. The energy transfer mechanism aims to efficiently harness wave energy from a depth below the water surface, where the water movement differs from that at the surface. This involves a column connected to a floating buoy on the wave surface, facilitating the conversion of wave movement from both vertical and horizontal directions primarily into the vertical direction. At the column’s end, multiple fins equipped with bimorph piezoelectric patches are strategically installed. The subsequent subsection delves into the specifics of designing the piezoelectric energy harvester transducer.

#### 2.3.1. Design of energy transfer mechanism

In this segment, the primary aim is to adeptly channel energy derived from surface waves, leveraging their substantial amplitudes, towards the deeper water regions characterized by reduced water current or diminished movement compared to the water surface. To simplify, the transition from the water surface to deeper levels results in a gradual decrease in water motion, a phenomenon visually depicted in [21-23].

As waves propagate through water, they don’t physically transport water like solid objects. Instead, they propagate energy through the water, causing water particles to undergo circular or elliptical motions. In deep water, these motions take a circular form. As a wave traverses, water particles within the wave execute back-and-forth orbits, without progressing along with the wave. The diameter of these circular orbits diminishes with depth, reaching a point where the circular motion becomes negligible [22, 23].

In shallow water, the wave interacts with the ocean floor, leading to flattened circular orbits of water particles, transforming into more elliptical shapes. The depth of the water significantly influences the trajectory of water particles. As depth increases, water particle motion gradually diminishes, reaching a depth where particles merely oscillate back and forth as the wave passes through.

In deep water, water particle trajectories follow a rotational path, with the rotational radius diminishing as one descends from the surface to deeper waters, indicating a reduction in motion. These regions with decreased motion are deemed secure zones for installing our piezoelectric transducer, specifically designed as articulated piezoelectric fins.

In this mechanism, the clamped section of the fin is firmly attached to a column rigidly linked to a buoy floating on the sea’s surface. This configuration effectively converts horizontal wave motion into vertical movement, as elucidated in Figure 7.

**Figure 7.**
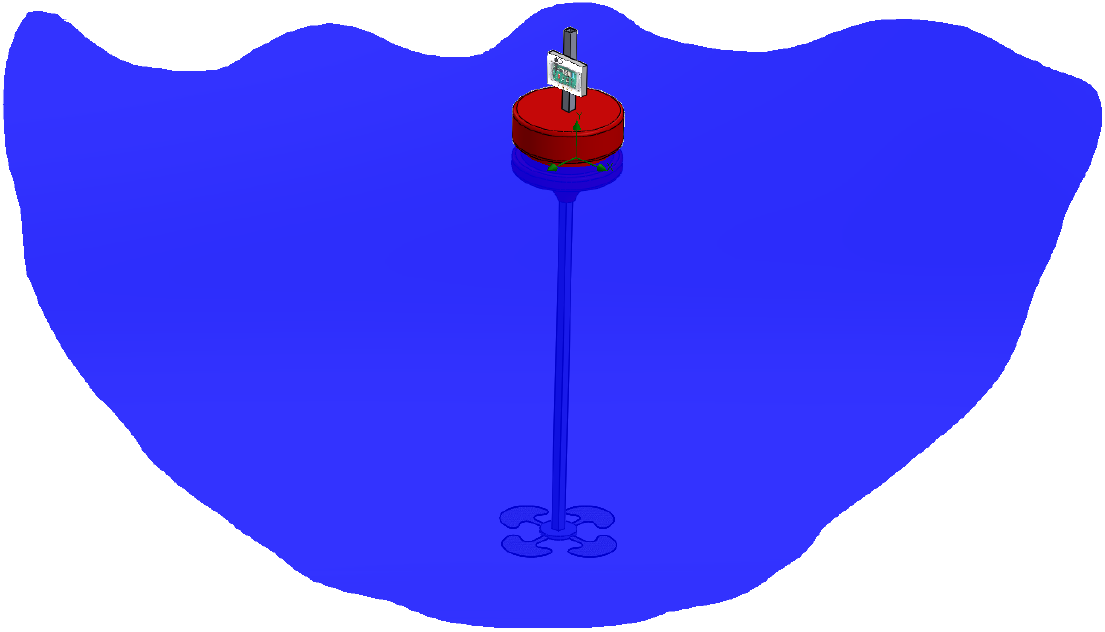
Schematic of sea wave energy transfer mechanism and installation of energy harvesters.

Figure 7 vividly outlines the operational configuration of this mechanism, featuring a buoy on the sea surface connected to a vertical column and multiple piezoelectric fins or transducers affixed at various levels along the column. The vertical movement of this column induces vibrations in the fins at an exceptionally low frequency, corresponding to the motion frequency of the column, inherently tied to the frequency and period of the sea waves.

#### 2.3.2. Design of piezoelectric energy harvester transducer

In this section, we delve into various experimental scenarios exploring different shapes and stiffness parameters for the transducer. The design of piezoelectric transducers is approached through two methods: harvesting energy either on off-resonance or on-resonance. The distinction lies in the natural frequency of the transducer, whether it falls outside or within the excitation frequency range from the ambient source, respectively. Given that the most likely excitation frequency from the ambient source is exceptionally low, below 1 Hz, designing a transducer for on-resonance application would necessitate a significantly larger size. Hence, the priority in this project is placed on designing a transducer for harvesting in an off-resonance range.

The chosen piezoelectric material for this case study is the DuraAct piezoelectric patch. This material proves suitable due to its composite flexibility and a Young’s modulus of 23.3 GPa. Additionally, being fully isolated makes it an ideal candidate for applications in sea water. As depicted in Figure 7, the configuration comprises four fins of identical shapes, securely clamped to the moving column connected to the buoy on the sea surface. Each fin undergoes morphing through two DuraAct patches positioned on the top and bottom of the substrate, consisting of a carbon fiber epoxy composite sheet with mechanical properties detailed in Table 3. The drawings and geometrical sizes of these composite sheets, along with prototypes of the piezoelectric fins, are showcased in Figure 8.

**Figure 8.**
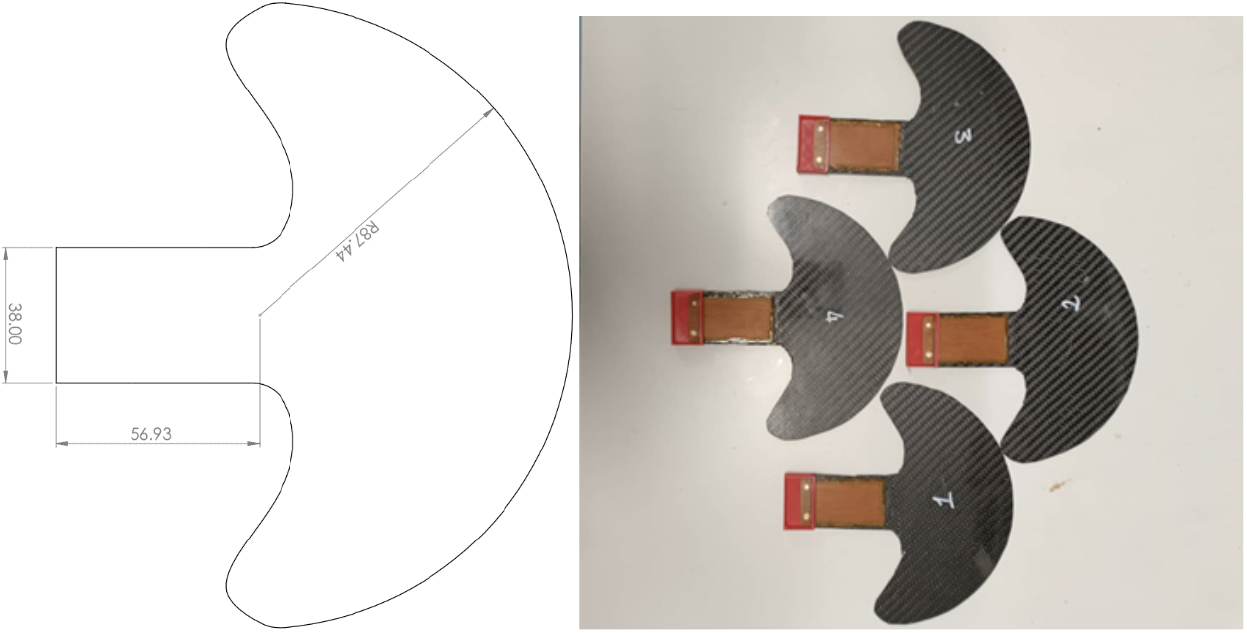
drawing of the substrate as a fin and four prototypes of the piezoelectric fins for the defined configuration.

This section provides an insight into the prototype and the experimental setup of the proposed sea wave piezoelectric energy harvester, as illustrated in Figure 9. In Figure 9.a, the prototype is showcased, featuring the configuration, geometry, and size of the piezoelectric fins along with the floating buoy on the water surface. The buoyancy line, crucial for calculating the distance between the fins’ surface and the water surface, is defined, and for this particular experimental setup, it measures approximately 1 meter. Notably, the depth of the water in the channel is 1.5 meters.

**Figure 9.**
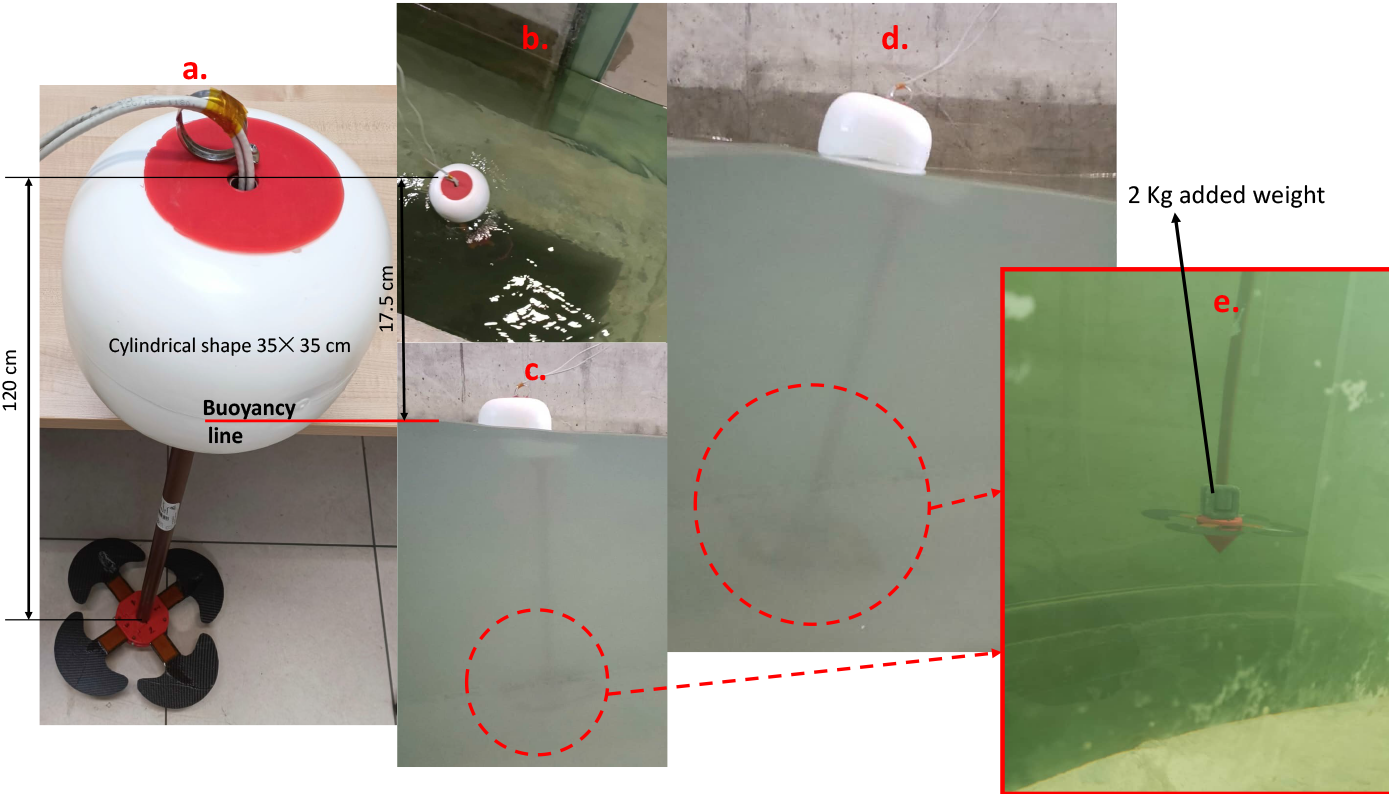
prototype and experimental setup of the piezoelectric energy harvesting from sea wave. a) prototype of the energy harvesting mechanism and defined buoyancy line, b) top view of the prototype in the water channel for test, c) side view of the prototype in water channel to show the real bouncy line, d) side view of the prototype in water channel to show the real bouncy line by passing surface wave, e) zoom in view of the fins and add mass for adjusting the buoyancy line in water channel.

Figures 9.b-d offer top and side views of the prototype within the water channel, vividly demonstrating the system’s stability when subjected to real water surface waves. Figure 9.e provides a closer view of the fins’ configuration, clamped to the connected column attached to the buoy on the water surface. In this depiction, a 2 kg weight is added to the end of the column to adjust the buoyancy line, ensuring it sits at a distance of 1 meter from the fins’ surface to the water free surface. The experiment takes place in a water channel with a depth of 1.5 meters, and the prototype is positioned at a considerable distance from the water wave generator in the channel. Surface waves reach the floating buoy when they attain a stable condition. This experimental setup is replicated for various scenarios involving surface waves of different heights and periods, as summarized in Figure 10.

**Figure 10.**
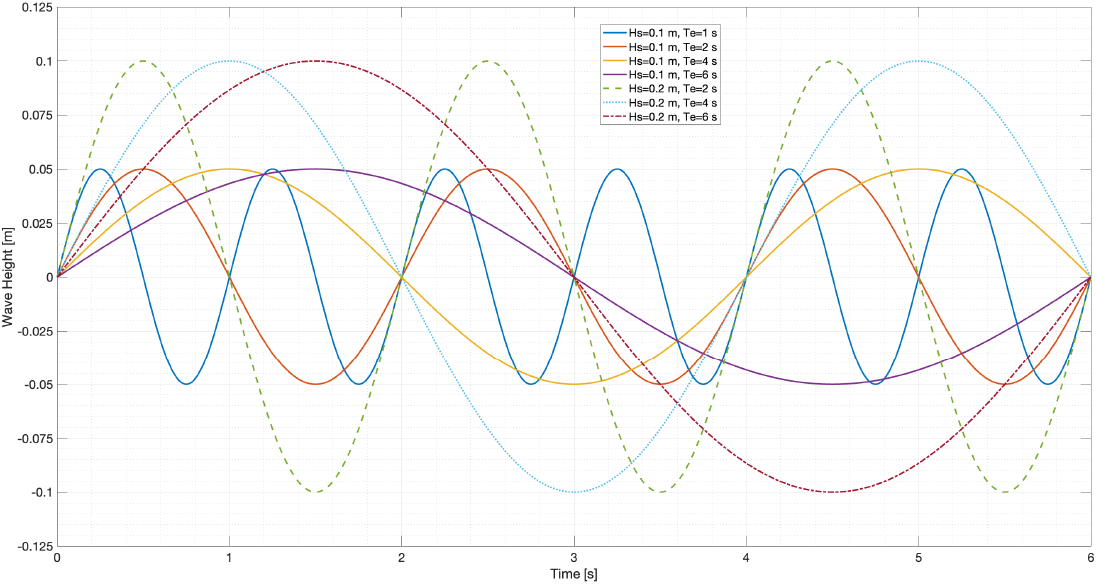
water surface wave passing pattern in different scenarios of wave heights and periods.

Figure 10 presents the surface wave patterns in various scenarios characterized by different wave heights and periods. The illustration features two amplitudes of wave heights—10 and 20 cm—across periods of 1, 2, 4, and 6 seconds. Notably, the figure zeroes in on the longest period, which is 6 seconds, offering a comparative analysis of surface wave patterns across several scenarios.

During this experiment, the surface wave pattern is consistently generated by the wave generator in the channel for a duration of 5 minutes. Measurements are conducted for 2 minutes within the middle of this wave generation, specifically when the surface wave shape attains stability as defined.

In this experimental setup, each of the eight piezoelectric patches, distributed across four fins, is connected to two wires for the measurement of the generated voltage through piezoelectric phenomena. This arrangement, comprising a total of 16 wires, provides the flexibility to inspect various electrical connections and introduce external loads between two bimorph patches for each fin. To ensure synchronized measurements for system identification through data analysis, a robust data logger with 8 differential multi-channels, the Analog and Digital Logger LGR-5329, has been selected.

The time history of generated voltages by each DuraAct patch, acting as a bimorph for each fin, is captured by the mentioned data logger in conjunction with a 1 MΩ resistor, as depicted in Figure 11. The measurement duration spans approximately 190 seconds, after which the total generated energy is calculated for each patch. Despite uniform materials and geometrical sizes of all patches and fins, variations in generated voltage and calculated energy arise due to manufacturing imperfections and the specific location of each fin during the passage of surface waves.

**Figure 11.**
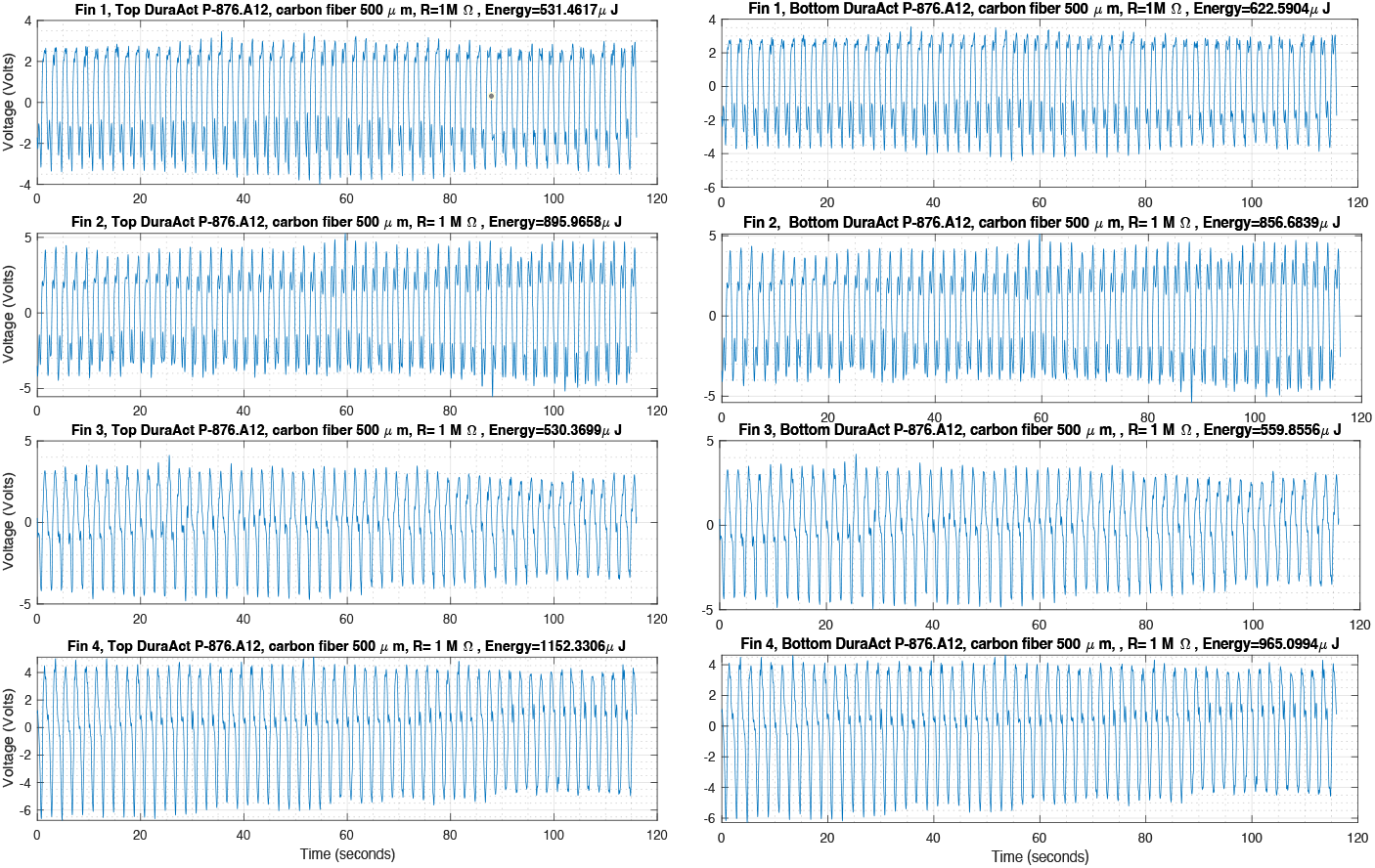
Time history of generated voltage and energy by each piezoelectric patch on the top and bottom side of each fin in parallel with 1 MΩ resistor as an external load and incident surface wave in 20 cm height and period 2 seconds.

In a specific test with a 20 cm surface wave height and a 2-second period, examining the plots in Figure 11 reveals that fins 2 and 4 outperform fins 1 and 3. Upon scrutinizing the configurations of the fins, it becomes evident that fins 2 and 4 are installed in opposite directions with a 180-degree difference, a configuration mirrored by fins 1 and 3. This performance disparity among fins is attributed to their distinct locations in the water during the passage of the surface wave.

The preceding test protocol is reiterated across various scenarios involving surface wave heights and periods, incorporating different external load values in parallel with the same fins. The outcomes of these tests are consolidated in Table 4, which facilitates a comparative analysis of four fins under different surface wave properties and parallel connections with external loads. The first three columns delineate seven distinct sea wave situations, varying in heights, periods, and the calculated wave power level (KWh/m), as determined by Eq. (1). Figure 12 visually supports the correlation between increasing wave height and period with elevated power levels.

**Table 4.** summary of the experimental results for comparing fins performance in several scenarios.

| Sea Wave |  |  | Fin 1 |  |  | Fin 2 |  |  | Fin 3 |  |  | Fin 4 |  |  |
| --- | --- | --- | --- | --- | --- | --- | --- | --- | --- | --- | --- | --- | --- | --- |
| Height (cm) | Period (s) | Power Level (KWh/m) | Generated Energy (mJ/min) |  |  | Generated Energy (mJ/min) |  |  | Generated Energy (mJ/min) |  |  | Generated Energy (mJ/min) |  |  |
| | | | 1 MΩ | 0.47 MΩ | 0.22 MΩ | 1 MΩ | 0.47 MΩ | 0.22 MΩ | 1 MΩ | 0.47 MΩ | 0.22 MΩ | 1 MΩ | 0.47 MΩ | 0.22 MΩ |
| 10 | 1 | 0.0049 | 10.5 | 9.4 | 5.8 | 30.8 | 27.8 | 17.3 | 60.3 | 54.4 | 33.8 | 31.6 | 28.5 | 17.7 |
| 10 | 2 | 0.0098 | 0.58 | 0.52 | 0.32 | 0.87 | 0.79 | 0.49 | 0.54 | 0.49 | 0.31 | 1.06 | 0.95 | 0.59 |
| 10 | 4 | 0.0197 | 0.028 | 0.026 | 0.016 | 0.029 | 0.027 | 0.017 | 0.024 | 0.022 | 0.014 | 0.028 | 0.025 | 0.016 |
| 10 | 6 | 0.0295 | 0.019 | 0.017 | 0.01 | 0.019 | 0.017 | 0.01 | 0.016 | 0.014 | 0.009 | 0.017 | 0.015 | 0.009 |
| 20 | 2 | 0.0393 | 4.38 | 3.95 | 2.45 | 4.28 | 3.86 | 2.4 | 10.33 | 9.31 | 5.78 | 7.034 | 6.34 | 3.94 |
| 20 | 4 | 0.0786 | 0.8 | 0.72 | 0.45 | 0.98 | 0.89 | 0.55 | 0.92 | 0.83 | 0.52 | 1.38 | 1.24 | 0.77 |
| 20 | 6 | 0.1180 | 0.52 | 0.47 | 0.29 | 0.71 | 0.64 | 0.39 | 0.47 | 0.42 | 0.26 | 0.53 | 0.47 | 0.29 |

**Figure 12.**
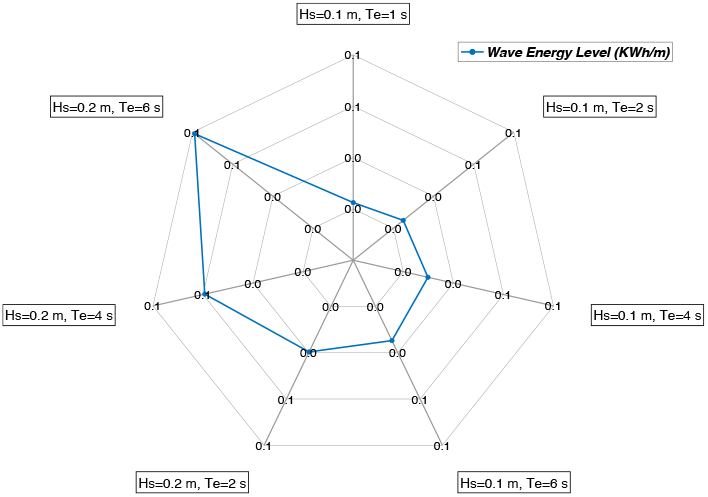
Radar plot of the wave power level in (KWh/m) versus wave heights and periods.

In the subsequent four main columns for each fin, there are three sub-columns, each listing the generated energy for the respective fin in parallel with the associated load. The energy scale in the table is denoted in mJ for one minute of measurement. Each row in the table is color-coded to correspond with the surface wave shape, detailed in Figure 10. A cursory glance reveals the piezoelectric energy harvesting system’s heightened sensitivity to the period or frequency of the surface wave. Despite the incremental increase in the generated energy with higher surface wave height, the system exhibits a more pronounced sensitivity to changes in wave period.

Figure 13 further elucidates the performance of the fins through three radar plots, showcasing the generated energy by the harvesting system under varying external load conditions. These figures highlight that the maximum harvested energy is attained with a 1 MΩ load in parallel with the fins, indicating an optimal external load for the piezoelectric transducer. Figures 13.a-c. illustrate a nearly linear decrease in harvested energy as the external load diminishes.

**Figure 13.**
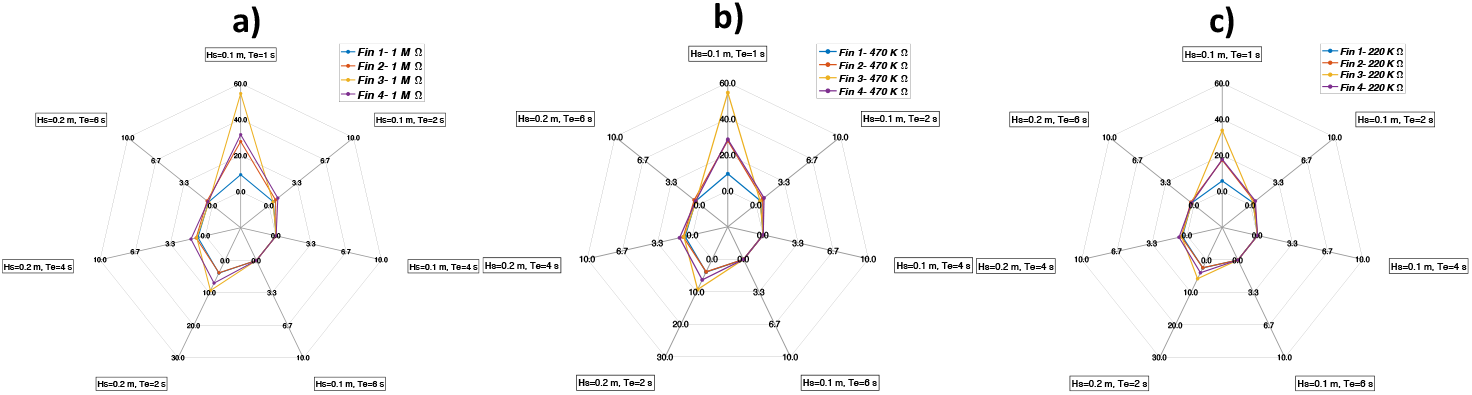
Radar plot of the generated energy by the designed energy harvester versus three different external loads. a) external load 1 MΩ, b) external load 470 KΩ and c) external load 220 KΩ

In anticipation and in alignment with real-world scenarios, especially drawing parallels with data collected from Maricoltura Mattinatese (41.725804, 16.109511) as depicted in Figures 4-6, this experiment is meticulously designed to mirror the most probable conditions—approximately 40% likelihood—of wave heights and periods observed in this marine location from 1993 to 2021. According to the joint probability distribution table in Figure 4, the most probable wave heights and periods, constituting around 18%, fall within the range of 10 cm to 20 cm and 1 second to 2 seconds, respectively. This specific range aligns with the design and testing parameters of the proposed piezoelectric energy harvesting system.

Data analysis and results discussion affirm that within this range, the proposed energy harvester exhibits optimal performance. In the best-case scenario, as per Table 4, one fin could harvest 60.3 mJ/min from a wave power level of 0.0049 KWh/m. In the second-ranked scenario, featuring a wave height of 20 cm, a period of 2 seconds, and a calculated power level of 0.0393 KWh/m, one fin could generate 10.33 mJ/min in the best case. A rough estimation, guided by Figure 5, suggests that within the most probable range of wave height, period, and power level at 0.0001 KWh/m, each fin of the harvester has the potential to generate at least 150 μJ/min in the worst-case scenario.

## 3. Conclusion

This paper introduces a pioneering piezoelectric energy harvester designed to convert energy from incident sea surface waves into electrical power, specifically tailored for self-powered IoT sensors in fish farming plants. The research journey commences with the collection of sea wave data spanning nearly 25 years, specific to the identified fish farming plant. This extensive dataset undergoes meticulous analysis, revealing critical insights such as joint probability distributions of surface wave height and period, sea wave power level, and normalized power level. These findings guide the identification of optimal points for designing an energy harvesting mechanism that aligns with the defined fish farming environment. The subsequent phases of the study involve the careful selection of suitable piezoelectric and substrate materials for constructing a piezoelectric fin. Finally, the energy harvesting mechanisms, encompassing both energy transfer and energy transduction systems, are intricately designed, fabricated, and subjected to testing in a controlled pool environment. The obtained results validate the feasibility of harvesting energy from sea surface waves, demonstrating the potential for generating dozens of mJ/min per each designed fin. As a next step, future work entails connecting multiple fins in parallel to a power conditioning circuit, aiming to charge the battery for the sensor system. This promising avenue represents the ongoing exploration in this field.

## Acknowledgements

This research was supported by fishRISE Project funded by Programma Operativo Nazionale “Ricerca e Innovazione”2014–2020 (PON”R&I”2014–2020) -”Remote, Intelligent & SustainableaquaculturE system for Fish (fishRISE)”n. ARS01_01053.”

## References

[1] Zambrano AF, Giraldo LF, Quimbayo J, Medina B, Castillo E. Machine learning for manually-measured water quality prediction in fish farming. PLOS ONE. 2021; 16(8):e0256380. 10.1371/journal.pone.0256380 PMID: 34407149.

[2] Boyd CE, Tucker CS. Pond Aquaculture Water Quality Management. 1st ed. Springer US; 1998. Available from: https://link.springer.com/book/10.1007/978-1-4615-5407-3.

[3] Mallya YJ, Thorarensen SPH. THE EFFECTS OF DISSOLVED OXYGEN ON FISH GROWTH IN AQUACULTURE. 2007.

[4] Antony AP, Lu J, Sweeney DJ. Seeds of Silicon: Internet of Things for Smallholder Agriculture; 2019. Available from: https://d-lab.mit.edu/resources/publications/seeds-silicon-internet-things-smallholderagriculture.

[5] Simbeye DS, Zhao J, Yang S. Design and deployment of wireless sensor networks for aquaculture monitoring and control based on virtual instruments. Computers and Electronics in Agriculture. 2014; 102:31–42. 10.1016/j.compag.2014.01.004.

[6] Smart Water Management IoT Solution | Water Quality Sensors Certified. Available from: https://www.libelium.com/iot-solutions/smart-water/.

[7] Medina JD, Arias A, Triana JM, Giraldo LF, Segura-Quijano F, Gonzalez-Mancera A, et al. (2022) Open-source low-cost design of a buoy for remote water quality monitoring in fish farming. PLoS ONE 17(6): e0270202. 10.1371/journal.pone.0270202.

[8] Shafiqur Rehman, Luai M. Alhems, Md. Mahbub Alam, Longjun Wang, Zakria Toor, A review of energy extraction from wind and ocean: Technologies, merits, efficiencies, and cost, Ocean Engineering 267 (2023) 113192. 10.1016/j.oceaneng.2022.113192.

[9] Kargar, S.M.; Hao, G. An Atlas of Piezoelectric Energy Harvesters in Oceanic Applications. Sensors 2022, 22, 1949. 10.3390/s22051949

[10] Renwen Liu, Lipeng He, Xuejin Liu, Shuangjian Wang, Limin Zhang, Guangming Cheng, A review of collecting ocean wave energy based on piezoelectric energy harvester, Sustainable Energy Technologies and Assessments 59 (2023) 103417. 10.1016/j.seta.2023.103417

[11] Shao-En Chen, Fu-Ting Pan, Ray-Yeng Yang, Chia-Che Wu, A multi-physics system integration and modeling method for piezoelectric wave energy harvester, Applied Energy 349 (2023) 121654. 10.1016/j.apenergy.2023.121654

[12] Shao-En Chen a, Wan-Yi Chen a, Ray-Yeng Yang b, Chia-Che Wu, A piezoelectric wave energy harvester equipped with a sequential-drive rotating mechanism and rotary piezoelectric harvesting component, Energy Conversion and Management: X 20 (2023) 100463. 10.1016/j.ecmx.2023.100463

[13] Xinru Du, Hidemi Mutsuda, Yoshikazu Tanaka, Takuji Nakashima, Taiga Kanehira, Naokazu Taniguchi, Yasuo Moriyama, Experimental and numerical studies on working parameter selections of a piezoelectric-painted-based ocean energy harvester attached to fish aggregating devices, Energy for Sustainable Development 71 (2022) 73–88. 10.1016/j.esd.2022.09.012

[14] Hakan Ucar, Patch-based piezoelectric energy harvesting on a marine boat exposed to wave-induced loads, Ocean Engineering 236 (2021) 109568. 10.1016/j.oceaneng.2021.109568

[15] Lipeng He, Renwen Liu, Xuejin Liu, Zheng Zhang, Limin Zhang, Guangming Cheng, A novel piezoelectric wave energy harvester based on cylindrical-conical buoy structure and magnetic coupling, Renewable Energy 210 (2023) 397–407. 10.1016/j.renene.2023.04.043

[16] Falnes, J., 2007. A review of wave-energy extraction. Mar. Struct. 20, 185–201.

[17] Goggins, J., Finnegan, W., 2014. Shape optimisation of floating wave energy converters for a specified wave energy spectrum. Renew. Energy 71, 208–220.

[18] Ali Azam, Ammar Ahmed, Hai Li, Alaeldin M. Tairab, Changyuan Jia, Ning Li, Zutao Zhang, Design and analysis of the optimal spinning top-shaped buoy for wave energy harvesting in low energy density seas for sustainable marine aquaculture, Ocean Engineering 255 (2022) 111434.

[19] Shadman, M., Estefen, S.F., Rodriguez, C.A., Nogueira, I.C.M., 2018. A geometrical optimization method applied to a heaving point absorber wave energy converter. Renew. Energy 115, 533–546.

[20] Yaakob, O., Hashim, F.E., Omar, K.M., Din, A.H.M., Koh, K.K., 2016. Satellite-based wave data and wave energy resource assessment for South China Sea. Renew. Energy 88, 359–371.

[21] http://www.thegeographeronline.net/coasts.html

[22] https://javalab.org/en/water_waves_en/

[23] https://www.thegeographeronline.net/coasts.html

